# Estimating evolutionary and demographic parameters via ARG-derived IBD

**DOI:** 10.1101/2024.03.07.583855

**Authors:** Zhendong Huang, Jerome Kelleher, Yao-ban Chan, David J. Balding

## Abstract

Inference of demographic and evolutionary parameters from a sample of genome sequences often proceeds by first inferring identical-by-descent (IBD) genome segments. By exploiting efficient data encoding based on the ancestral recombination graph (ARG), we obtain three major advantages over current approaches: (i) no need to impose a length threshold on IBD segments, (ii) IBD can be defined without the hard-to-verify requirement of no recombination, and (iii) computation time can be reduced with little loss of statistical efficiency using only the IBD segments from a set of sequence pairs that scales linearly with sample size. We first demonstrate powerful inferences when true IBD information is available from simulated data. For IBD inferred from real data, we propose an approximate Bayesian computation inference algorithm and use it to show that poorly-inferred short IBD segments can improve estimation precision. We show estimation precision similar to a previously-published estimator despite a 4 000-fold reduction in data used for inference. Computational cost limits model complexity in our approach, but we are able to incorporate unknown nuisance parameters and model misspecification, still finding improved parameter inference.

**Author summary:** Samples of genome sequences can be informative about the history of the population from which they were drawn, and about mutation and other processes that led to the observed sequences. However, obtaining reliable inferences is challenging, because of the complexity of the underlying processes and the large amounts of sequence data that are often now available. A common approach to simplifying the data is to use only genome segments that are very similar between two sequences, called identical-by-descent (IBD). The longer the IBD segment the more informative about recent shared ancestry, and current approaches restrict attention to IBD segments above a length threshold. We instead are able to use IBD segments of any length, allowing us to extract much more information from the sequence data. To reduce the computation burden we identify subsets of the available sequence pairs that lead to little information loss. Our approach exploits recent advances in inferring aspects of the ancestral recombination graph (ARG) underlying the sample of sequences. Computational cost still limits the size and complexity of problems our method can handle, but where feasible we obtain dramatic improvements in the power of inferences.

## Introduction

A common data-reduction technique when analysing samples of genome sequences is to identify identical-by-descent (IBD) genome segments [1–5]. In practice IBD is often identified by searching for regions with no evidence for recombination along two sequences since their most recent common ancestor (MRCA). Further, only IBD segments (IBDs) above a given length threshold, often 2 to 4 cM, are retained. This practice wastes valuable information, but has been necessary because the inference of short IBDs is too noisy to be useful for downstream analyses.

The ancestral recombination graph (ARG) is widely used to represent the genealogical history of a sample [6–8] and recent developments in inferring aspects of the ARG [9–13] now permit us to rapidly extract IBD directly from inferred shared ancestors, without requiring zero recombination. Further, computationally fast ARG inference and extraction of IBD can be implemented within an approximate Bayesian computation (ABC) algorithm which removes the need for an information-wasteful length threshold. Instead, we reduce computational cost by using an efficient subset of IBDs that scales linearly with sample size with little information loss relative to using all IBDs.

Our approach relies on an efficient data structure encoding features of an ARG underlying a sample of genome sequences, called the succinct tree sequence (TS) [14]. The TS minimises redundant storage of subsequences that are similar due to shared ancestry. It has led to spectacular improvements in storage and simulation of large genome datasets [15], and has recently been applied to IBD-based inferences about demographic history and evolutionary parameters [16].

We first demonstrate powerful inferences of mutation and sequencing error rates, TMRCA (time since the MRCA), and past and present population sizes, given true IBD information in simulation studies. For real datasets, we propose TSABC: ABC with statistics computed from IBDs extracted from an inferred TS. We demonstrate the performance of TSABC with inferences of the mutation rate and population size in simulation studies and real data, and we compare mutation rate estimates with previously-published results and with analyses using a range of IBD length thresholds.

We find that using IBDs extracted from an inferred ARG leads to a surprisingly small loss of precision relative to use of true IBDs. Further, even a low threshold on IBD length reduces the quality of inferences, despite the fact that short IBDs are poorly inferred. TSABC is computationally demanding, which limits the size and complexity of inference problems that can be tackled. However, TSABC can achieve comparable results to previous estimators using much smaller data sets: we show similar precision to a previously-published estimator despite a 4 000 fold reduction in data available for inference.

## Methods

### Definition and notations

The TS encodes genome sequence data efficiently by storing subsequences that are similar due to shared ancestry as variations of an ancestral sequence. It is defined [17] as, *{ C, P, E, ℳ}*, where C=*{* 1,…, *m}* is the set of leaf (or tip) nodes corresponding to *m* observed sequences each of length *l*, and *P* =*{m*+1,…, *n}* is the set of internal (ancestral) nodes of the TS ordered backwards in time from the present. An edge in *E* = *{*(*c*_*i*_, *p*_*i*_, *l*_*i*_, *r*_*i*_): *i* = 1, 2,…, *I}* represents inheritance of sites in the segment [*l*_*i*_, *r*_*i*_], with 1 *≤ l*_*i*_ *≤ r*_*i*_ *≤ *l**, from internal node *p*_*i*_ *∈ P* to its child *c*_*i*_ *∈ {*1,…, *p*_*i*_*−*1*}*, while ℳ = *{*(*c*_*j*_, *s*_*j*_): *j* = 1, 2,…*}* stores the set of sites *s*_*j*_ at which there is a sequence difference between *c*_*j*_ and its parent, due either to a mutation or, if *c*_*j*_ is a leaf node, sequencing error. The TS has the “succinct” property that any tree component conserved over a genome segment is stored only once, which greatly reduces data storage requirements compared with retaining all distinct marginal trees.

### Identity by descent and efficient subsets

We denote the *i*th IBD segment in the TS by IBD_*i*_ = (*c*_*i*1_, *c*_*i*2_, *l*_*i*_, *r*_*i*_, *p*_*i*_, *M*_*i*_), *i* = 1,…, *I*, ordered such that *c*_*i*1_ is non-decreasing in *i*. Here *c*_*i*1_ and *c*_*i*2_ are the leaf nodes of the two sequences, [*l*_*i*_, *r*_*i*_] is the IBD genome segment, *p*_*i*_ is the MRCA node of *c*_*i*1_ and *c*_*i*2_ for this segment, and *M*_*i*_ denotes the set of sites in [*l*_*i*_, *r*_*i*_] at which *c*_*i*1_ and *c*_*i*2_ differ. As there is no length threshold, the IBDs of any sequence pair partition the genome: every sequence site is included in exactly one of the IBD segments.

Each IBD_*i*_ has the same MRCA at each site in [*l*_*i*_, *r*_*i*_], and a different MRCA at adjacent sites. Imposing a no-recombination requirement as part of the definition of IBD would be more restrictive, since the absence of recombination implies a common MRCA but the reverse does not hold (see Figure 1, left, for examples of recombinations that do not change the MRCA).

**Fig 1.**
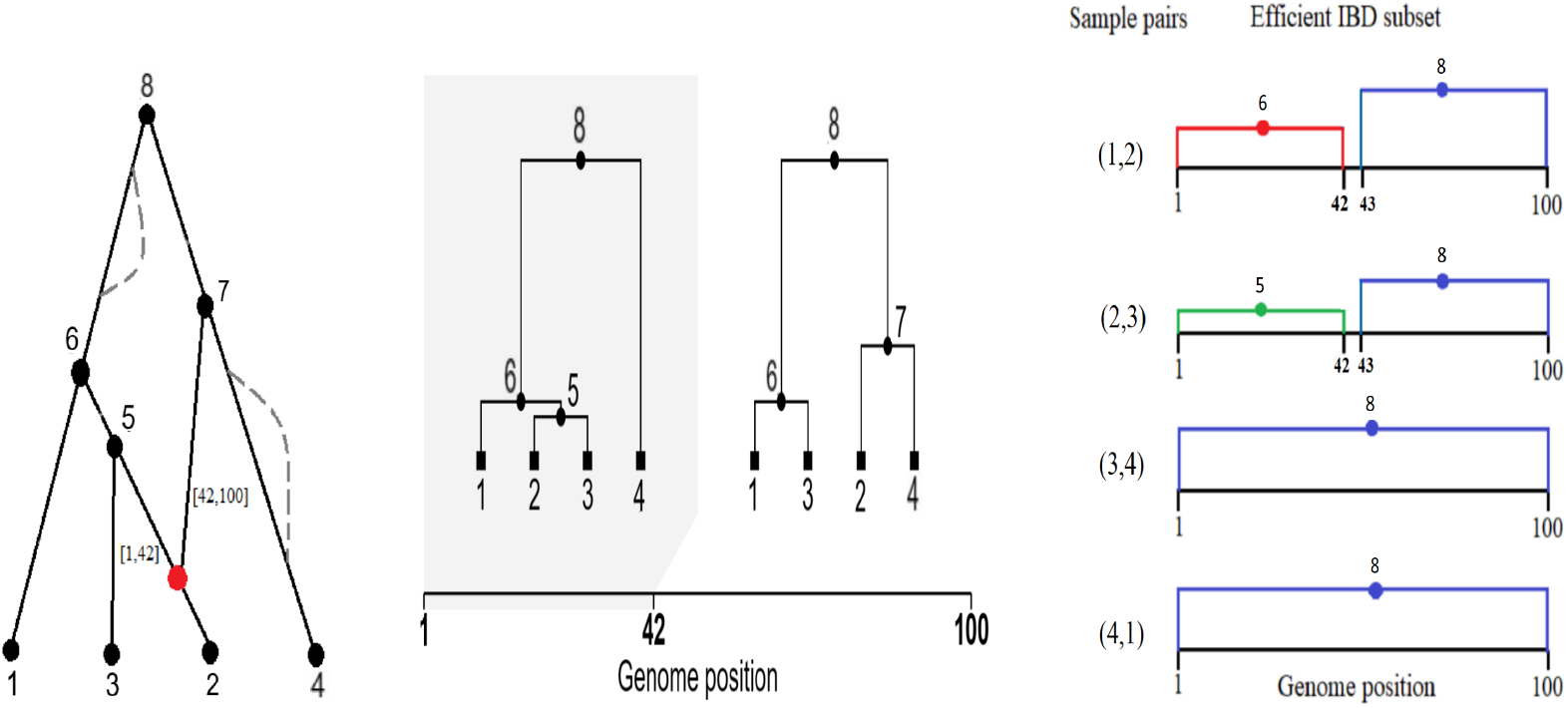
An ancestral recombination graph (ARG) spanning a genome sequence of length **l** = 100 (left), the corresponding sequence of local trees (middle) and efficient IBD subset (right). The ARG has leaf nodes *{* 1, 2, 3, 4*}* = C, named ancestral nodes *{*5, 6, 7, 8*}* = *P*, and a recombination at site 42 of an internal ancestral node (red dot). The two dashed lines in the ARG represent inheritance paths due to two ineffective recombination events, which are not represented in the TS. The efficient IBD subset includes two IBD segments for the node pair (1, 2), corresponding to intervals [1, 42] and [43, 100] which have MRCA 6 and 8, respectively, and one IBD segment spanning the whole sequence for pairs (3, 4) and (4, 1).

To reduce computational effort, we use for inference only an “efficient” subset of IBDs. After fixing an arbitrary order for the sequences, we include in the subset only the IBDs of the sequence pairs (1, *m*) and (*c, c*+1) for *c* = 1,…, *m−*1 (see Figure 1, right, and Appendix S1). An efficient subset has the property that each edge of the TS is included in a descent path from the MRCA for at least one IBD segment in the subset, which ensures that information is retained in the subset about every mutation.

Imposing a length threshold on IBDs is also a form of data reduction but we show below that it can lead to high information loss, because mutations are ignored if they occur at sites not contained in a sufficiently long IBD segment.

### Estimation

Let *µ* and *ϵ* be the per-site per-generation mutation rate and the per site sequencing error rate, both assumed constant over sites. For *i* = 1,…, *I*, let *g*_*i*_ denote the age of *p*_*i*_ in generations, and let *N* (*g*), *g* = 0, 1, 2,…, be the population size *g* generations in the past. In Appendix S2 we derive method-of-moment estimators for *µ* and *ϵ*, and non-parametric estimators of *g*_*i*_, *i* = 1,…, *I*, and *N* (*g*), *g* = 0, 1, 2,…, based on statistics computed from IBD lengths. We investigate the performance of these estimators when true IBD information is available in simulation studies. The recombination rate *r* is assumed constant over sites and known for all inferences; the extension to a known recombination map is straightforward (a recombination at site *s* means between sites *s* and *s* + 1).

For observed sequence data, true IBD information is not available and we extract IBDs from an inferred TS. TSABC uses summary statistics derived from these IBDs and related to the method-of-moments estimators. For inference of *µ* and *ϵ*, we use the Statistics 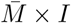 and *C*_1_ (Appendix S2.1) which are linear transformations of the method-of-moments estimators 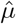 and 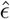. Nonparametric estimation of *N* (*g*) is not feasible, but we can estimate the parameters of a demographic model, which allows powerful inference provided that the model is adequate. We use as statistics the mean and standard deviation (SD) of IBD lengths *r*_*i*_ *− l*_*i*_, *i* = 1,…, *I*.

### Simulation study design: true IBD available

We jointly estimated *µ, ϵ, g*_*i*_, *i* = 1,…, *I*, and *N* (*g*), *g ≥* 0, using our novel estimators. We used msprime [19] to generate TS under the coalescent with recombination model [20, 21], assuming demographic models C, Ga and S (Table 1). From each TS we extracted an efficient subset of IBDs (Algorithm 1). Sequencing error was simulated by adding elements to ℳ at leaf nodes of the generated TS. At the largest error rate (*ϵ* = 10^*−*3^), any singleton variant is a few times more likely to arise from sequencing error rather than a mutation.

**Table 1.** Parameter values, sample properties and demographic models for the simulation study. Unless otherwise stated, 25 simulation replicates were generated in each scenario. Model Ga is used for inferences given true IBD and Model Gb is used for inferences from inferred IBD. The value of *r* is assumed known for all inferences, whereas *µ, ϵ* and *N* (*g*), *g ≥* 0, are targets of inference.

| Evolutionary parameters and sample properties |  |  |
| --- | --- | --- |
| Symbol | Definition | Value(s) in simulations |
| $\mu$ | mutation rate | $1.3 \times 10^{-8}$ per site per generation |
| $\epsilon$ | sequencing error rate | up to $10^{-3}$ per site |
| $r$ | recombination rate | $10^{-8}$ per site per generation |
| $m$ | sample size | 10, 20, 40, 60, 80, 100, 160 or 200 sequences |
| $\ell$ | sequence length | $10^6$, $10^7$ or $10^8$ sites |
| Demographic models ($N(g)$ = population size $g$ generations ago) | | |
| Model C | $N(g)$ constant | $N(g) = 2 \times 10^4$ |
| Model G | $N(g) = N(0) \times e^{-\tau g}$ | $N(0) = 10^6$, $\tau = 10^{-4}$ (Model Ga)<br>$N(0) = 2 \times 10^5$, $\tau = 10^{-3}$ (Model Gb) |
| Model S | $N(g) = N(0)$ for $0 < g < G$<br>$= N(G)$ for $g \geq G$ | $N(0) = G = 4 \times 10^4$<br>$N(G) = 10^4$ |
| Model EA | European-American demographic history [18] | See Figure S2 for $N(g)$ values;<br>gene conversion: tract length 100 bp<br>rate $10^{-8}$/site/generation. |

### Simulation study design: inferred IBD

We used msprime to generate simulated sequences, recoded them as binary strings using 0 and 1 for the ancestral and derived alleles and added sequencing errors by assigning 1 to randomly selected sites at rate *ϵ* (see [22] for alternative models of sequencing error). We choose tsinfer [10] to infer the TS from the resulting sequence data; speed is critical for an ABC algorithm, and tsinfer is the fastest of the current methods, while retaining high accuracy [13, 23]. Unless otherwise stated, in each scenario we used *η* = 2 500 simulations with ABC acceptance rate 0.05 (125 acceptances).

We first use simulations to confirm previous reports [24] that the quality of IBD inference is often poor, particularly for short IBDs. We compared the number of true and inferred IBDs for datasets simulated under Model C with *µ* ranging from 1 to 20 units of 10^*−*8^ per site per generation and *m* = 10, 20 and 160. We also compared the length distribution of true and inferred IBDs for *m* = 160 and *µ* = 1.3 *×* 10^*−*8^.

To investigate the effect of including short IBDs, both true and inferred, we also modified TSABC to include only IBDs with length greater than a threshold of 1, 2 or 4 units of 10^4^ bp. When a threshold was applied, we included all IBDs satisfying the threshold, rather than using only the efficient subset of IBDs.

We next investigated TSABC estimation of *µ* under Model C and Model Gb with **l** = 10^7^. The *N* (*g*) values and *ϵ* = 0 were assumed known for the inference and we adopted a Uniform(10^*−*8^, 2 *×* 10^*−*8^) prior distribution for *µ*. For the Model C simulations with *m* = 10, we also applied TSABC after thresholding on IBD length and repeated using true IBD extracted from the msprime simulations.

To study TSABC estimation of the population size *N* (*g*), we used *m* = 200 and **l** = 10^6^ under each of Model C and Model Gb. For both data simulation models, the TSABC inference used Model G but with different prior distributions. When the simulation model was Model C, we fitted Model G with independent prior distributions Uniform(10^4^, 3 *×* 10^4^) for *N* (0) and Uniform(*−*2 *×* 10^*−*5^, 2*×* 10^*−*5^) for *τ*. Whenever *τ <* 0, we impose a population size limit *N* (*g*) *≤* 2 *N ×* (0). With simulation model Model Gb, the independent prior distributions were Uniform(10^5^, 3 *×* 10^5^) for *N* (0), and Uniform(0, 0.002) for *τ*. All parameters were treated as known except the targets of inference *N* (0) and *τ*.

We performed additional simulations to allow comparison with the inferences of *µ* reported by [18]. Data were simulated under Model EA, which aims to capture key features of the demographic history of European-Americans (Table 1), and Model C modified to include sequencing errors. The Model C simulations of [18] used *ϵ* = 10^*−*4^ but no gene conversion, while for Model EA they set *ϵ* = 0 and included gene conversion. We include both sequencing error and gene conversion in both Model C and Model EA simulations. We used a 400-fold smaller sample size than [18] (*m* = 10 versus *m* = 4 *×* 10^3^) and 10-fold smaller genome length (**l** = 10^7^ per chromosome, versus **l** = 10^8^).

We did not include gene conversion in the TSABC inference, thus challenging it with model misspecification. As a further challenge, we treated *N* (*g*) as unknown when inferring *µ*, and misspecified the model for *N* (*g*) in the TSABC simulations.

When the data were simulated under Model C, TSABC used independent prior distributions Uniform(10^*−*8^, 2 *×* 10^*−*8^) for *µ* and Uniform(0.6 *×* 10^*−*4^, 1.6 *×* 10^*−*4^) for *ϵ*. For *N* (*g*), we adopted Model G with independent priors *N* (0) *∼* Uniform(14 000, 30 000) and *τ ∼* Uniform(*−*2 *×* 10^*−*5^, 10^*−*5^).

When the data were simulated under Model EA, TSABC used a Uniform(10^*−*8^, 2 *×* 10^*−*8^) prior distribution for *µ*. For inference of *N* (*g*), we adopted Model S with independent prior distributions *N* (0) *∼* Uniform(11 000, 15 000), *G ∼* Uniform(4500, 6500) and *N* (*G*) *∼* Uniform(45 000, 49 000).

### Mutation and growth rates in the 1000 Genome Project

We analyse chromosomes 20 and 21 from 8 of the 26 human populations of the 1000 Genomes Project (1KGP) [25] making use of the demographic model of [26] which we refer to as the 1KGP model. See Figure S2 for plots of the 1KGP model and Appendix S3 for details of the data analysis. Separately for each chromosome, we use TSABC to infer *µ* assuming the prior Uniform(10^*−*8^, 2 *×* 10^*−*8^) and the 1KGP model. The 16 sets of 125 accepted values were analysed in a two-way ANOVA to assess differences in *µ* across chromosomes and over populations.

Next, we use chromosome 20 and 21 data to estimate population size *N* (*g*) assuming the 1KGP model for *g ≥* 1000 and fitting demographic Model G for 0 *≤ g ≤* 1 000, constrained such that *N* (1000) in Model G matches the 1KGP model value. The constrained Model G has one free parameter *N* (0), for which we adopt a Uniform(10000, 240000) prior distribution. To reduce computational effort with little loss of information, in both the observed dataset and TSABC simulations we removed SNPs with a minor allele count *>* 40, which typically arose at *g ≫* 1 000. We estimate *N* (*g*) from each chromosome separately and average the results.

## Results

### Simulation study results: true IBD available

While use of the efficient subset of IBDs reduces computational cost in proportion to the reduction in sequence pairs from *m*(*m−* 1)*/*2 to *m*, the average estimated SD of 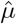 in our simulation study increased only slightly, from 0.017 to 0.019 units of 10^*−*8^ (see also Figure S3, left panel). This gain in computation time is typically worth the small loss of statistical efficiency.

Both 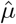 and 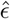 are well estimated in all demographic models, with no indication of bias (Figure 2). Increasing *m* has only a modest effect on the SD of estimators, whereas **l** has a larger effect (SD scales with 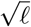, Figures 2, S3 (right) and S4). Sequencing errors only inflate the number of singleton variants, so 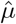 is little affected by increasing *ϵ* (Figure 2).

**Fig 2.**
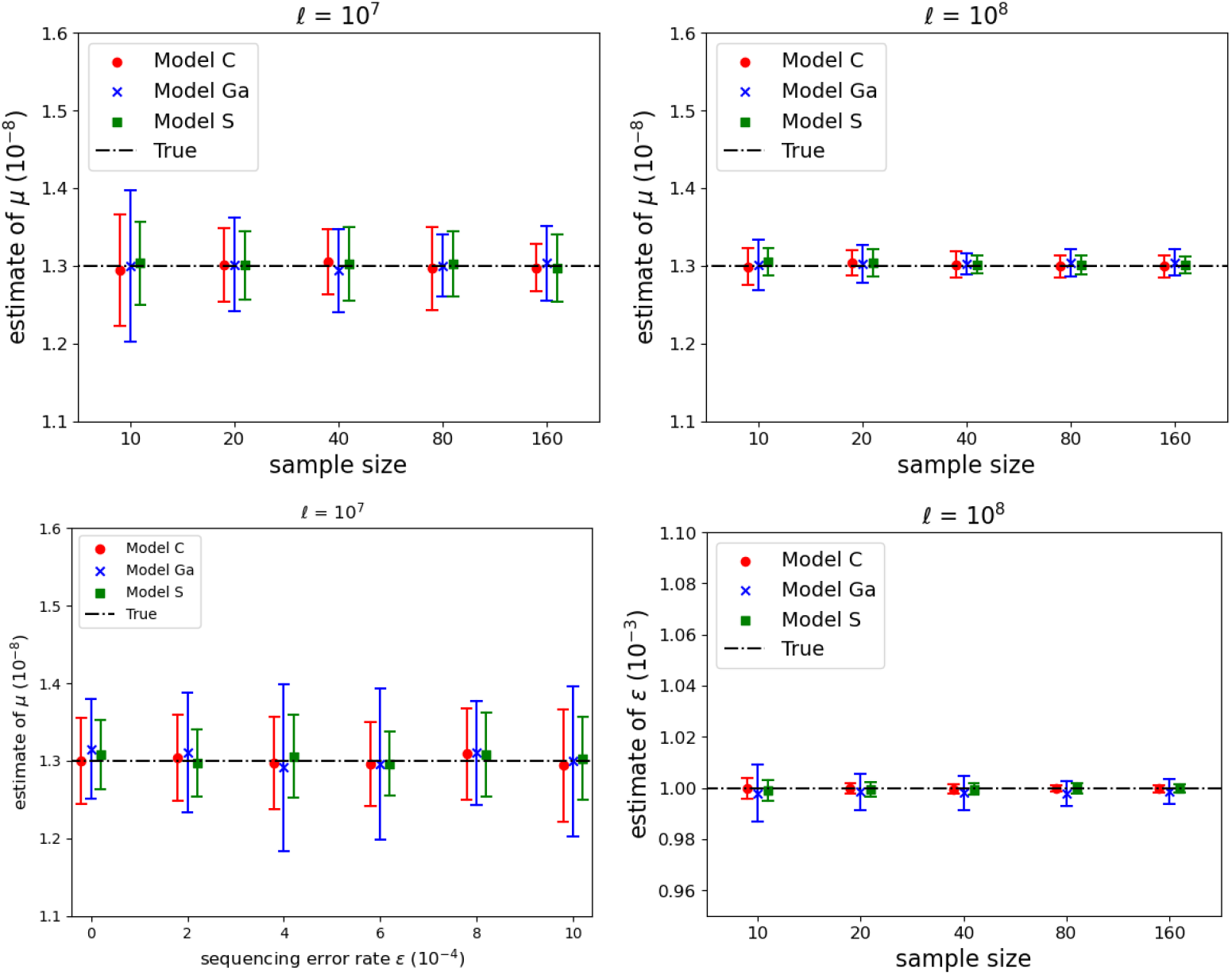
Inference of mutation rate *µ* and sequencing error rate *ϵ* with two sequence lengths (columns), when true IBD was available for inference. Line segments show indicative 95% CIs computed from the average estimate (indicated by a symbol, see legend box) and the empirical SD of the estimates from 25 simulated datasets in each scenario. Bottom left panel shows the impact of *ϵ* On 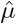 when *m* = 10, in the other three panels *ϵ* = 10^*−*4^.

Although individual 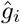 are not precise, the empirical and theoretical densities obtained from all 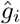, *i* = 1…, *I*, are close (Figure 3) despite the TS used for input only including information about the order of the coalescent events, and not their times. The population size estimator 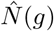 is accurate under all models, at least for *g ≤* 5 *×* 10^5^ (Figure 4). Figures S3 (right) and S4 show more precise estimates of 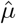 and *N* (*g*), with a longer sequence length.

**Fig 3.**
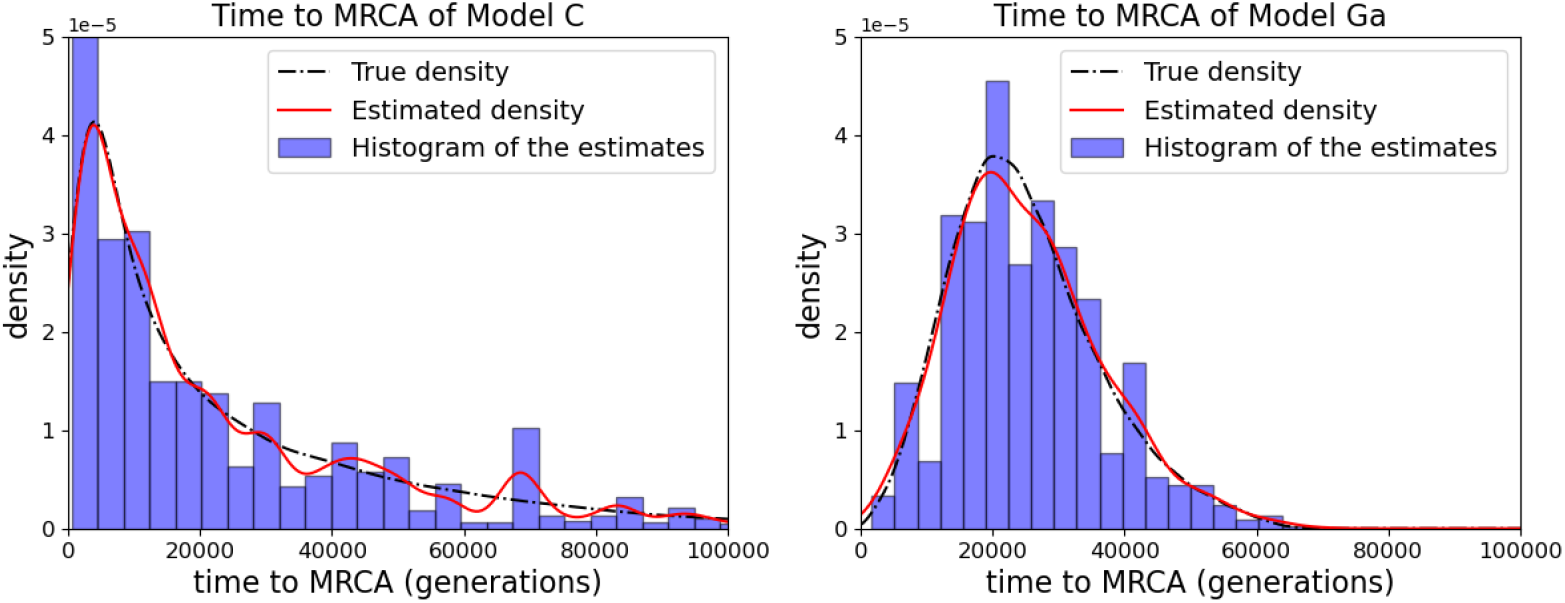
Histogram of the 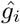, *i* = 1,…, *I*, obtained from one sample simulated under each of Model C (left) and Model Ga (right), with sample size *m* = 80, sequence length **l** = 10^8^ and sequencing error rate *ϵ* = 10^*−*3^. Also shown is a probability density obtained by kernel smoothing of the 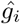 together with the true density. True IBD was available for inference but no time information.

**Fig 4.**
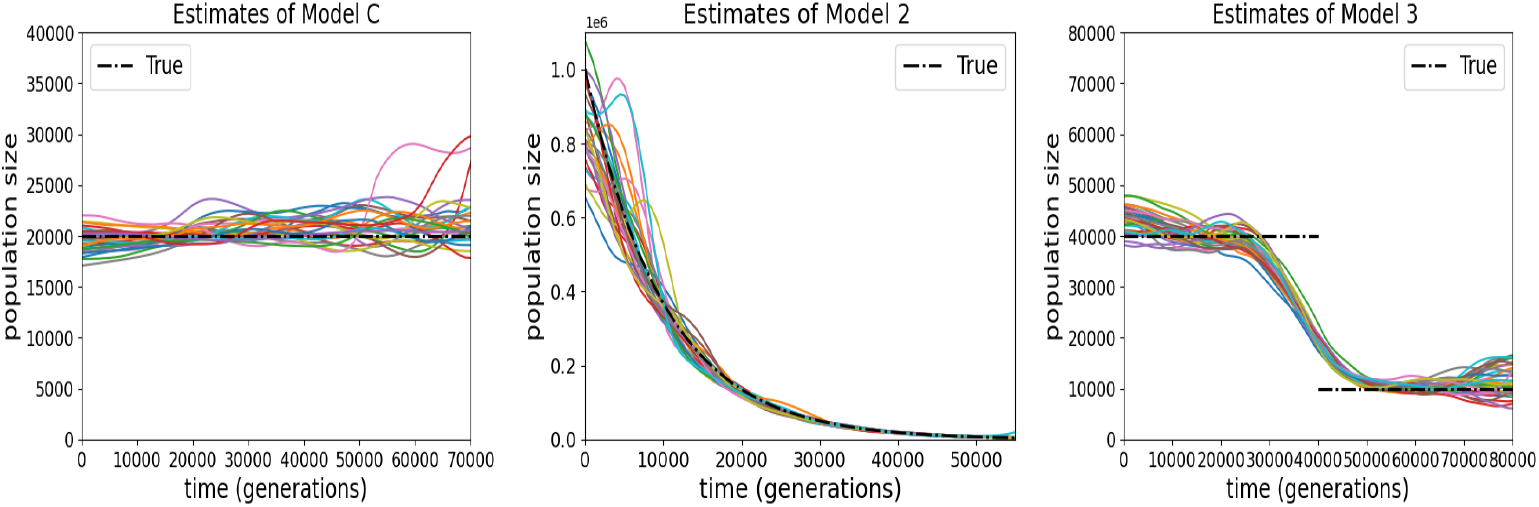
Estimates of the population size *N* (*g*), *g ≥* 0, from each of 25 simulation replicates under Model C, Model Ga and Model S, when true IBD was available for inference. Sequence length is **l** = 10^8^, sequencing error rate is *ϵ* = 10^*−*3^ and sample size is *m* = 80.

### Simulation study results: inferred IBD

The number of inferred IBDs tends to increase with both *µ* and *m*, but except for very high *µ* (over 10 times the average human value when *m* = 160) it remains well below the true number of IBDs (Figure 5, left). Correspondingly, the length distribution of inferred IBDs is highly skewed towards larger values relative to the true distribution (Figure 5, right), as previously reported [27, 28]. Despite this poor detection of small IBDs, and consequent tendency for inferred IBDs to be longer than the true IBDs, Table 2 shows that each increase in the length threshold reduced the precision of inference, both for inferred and true IBD, so that even poorly-inferred short IBDs do contribute useful information for inference. We also see in Table 2 (final column) further evidence that use of the efficient subset of IBDs leads to only a small loss of statistical efficiency. As expected, the use of true IBD improves TSABC compared with using inferred IBD, but the magnitude of the improvement is modest in the case of standard

**Table 2.** Comparison of TSABC inference for *µ* using different IBD length thresholds. Each result is an average over 25 Model C simulation replicates with *m* = 10 and *ϵ* = 0. In the last column, values based only on IBDs in the efficient subset are given in ().

| Threshold ($10^4$ bp) | 4 | 2 | 1 | 0 | |
| --- | --- | --- | --- | --- | --- |
| inferred IBD |  |  |  |  |  |
| # IBD | 1 001 | 7 033 | 24 683 | 48 289 | (10 667) |
| $\hat{\mu}$ ($10^{-8}$) | 1.51 | 1.44 | 1.32 | 1.30 | (1.31) |
| SD ($10^{-8}$) | 0.219 | 0.167 | 0.067 | 0.041 | (0.043) |
| true IBD |  |  |  |  |  |
| # IBD | 478 | 1 803 | 6 940 | 121 027 | (26 394) |
| $\hat{\mu}$ ($10^{-8}$) | 1.33 | 1.30 | 1.31 | 1.30 | (1.29) |
| SD ($10^{-8}$) | 0.141 | 0.089 | 0.064 | 0.029 | (0.034) |

**Fig 5.**
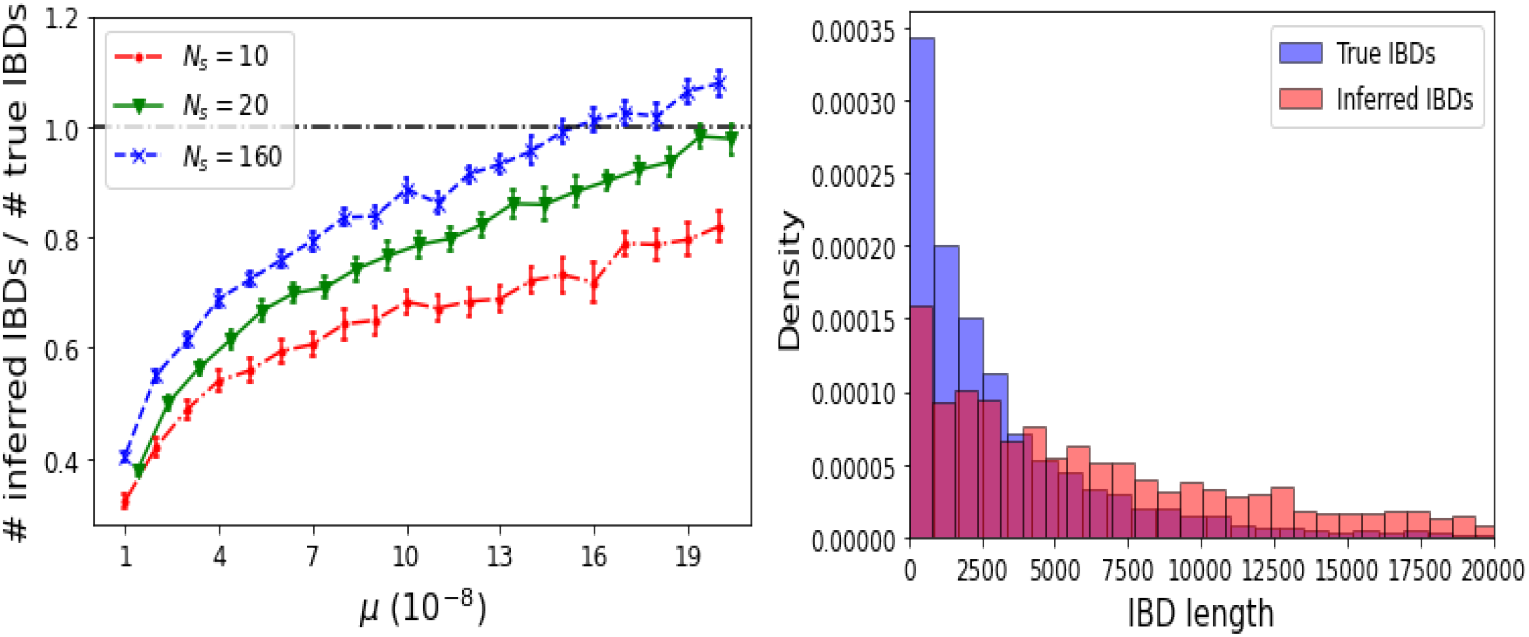
Comparison of true and inferred IBDs. Left: each symbol and vertical line segment shows the mean and 95% CI of the mean ratio of IBD counts over 25 Model C simulations with sample sizes *m* = 10, 20 and 160. Right: histograms of true and inferred IBD length distributions for a Model C simulated dataset with *m* = 160 and sequence length **l** = 10^6^.

TSABC (threshold = 0). For higher thresholds, bias can be high due to low precision of inference and the prior boundary at 10^*−*8^.

Although TSABC can provide approximations to the full posterior distribution, we report here only posterior mean estimates of unknown parameters. For inference of *µ*, it appears that any bias of TSABC is small for both models (Figure 6). Some under-estimation is expected because the binarisation of the sequence data obscures instances of multiple mutations at the same site, but this effect is negligible.

**Fig 6.**
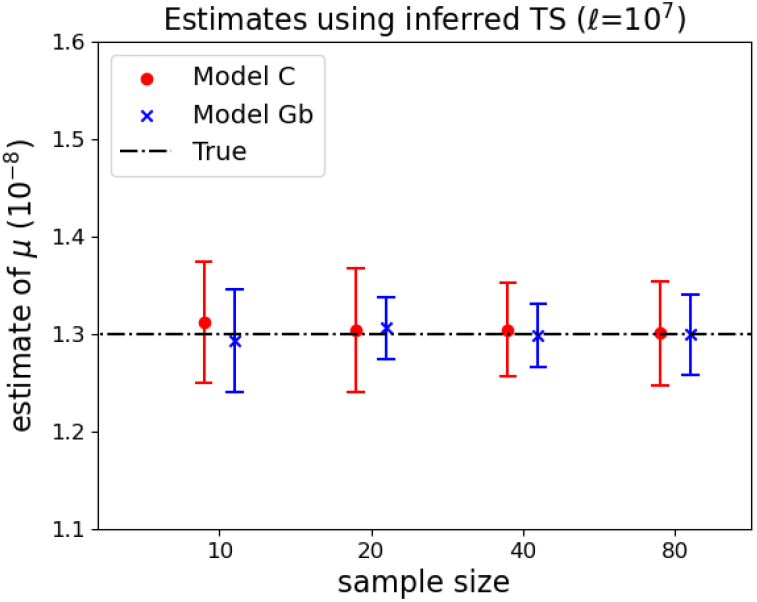
TSABC estimation of mutation rate *µ*. Symbols and line segments show mean and 95% CI over 25 simulations with no sequencing error (*ϵ* = 0) and sequence length **l** = 10^7^.

Parametric estimation of *N* (*g*) also performs well (Figure 7). When the data simulation model was Model C, the average estimate of *N* (0) (true value 20 000) over the 25 replicates is 20 931 with standard error (SE) 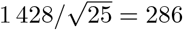,while e for the growth rate *τ* (true value 0) the average estimate *±* SE is 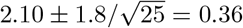 (in units of 10^*−*6^). When the data simulation model was Model Gb, for *N* (0) (true value 200 000) we obtained 202 534 *±* 2 173 while for *τ* (true value 1) we obtained 1.08 *±* 0.07 (in units of 10^*−*3^).

**Fig 7.**
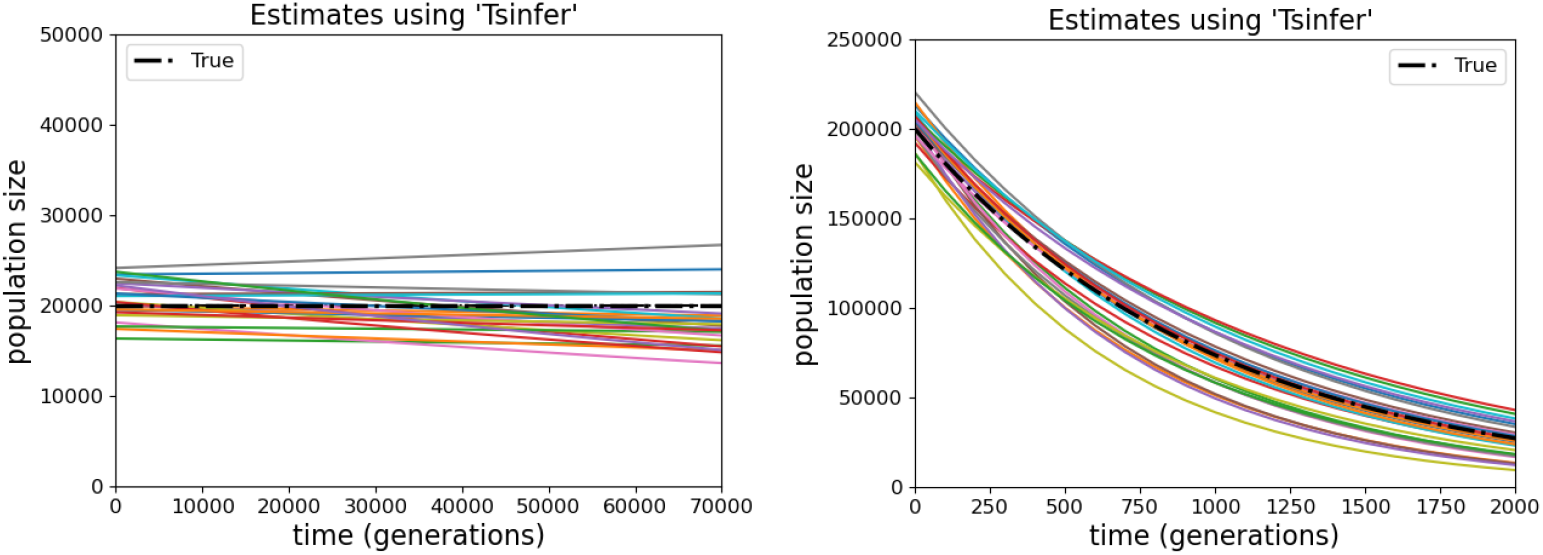
Fitted exponential curves for the population size *N* (*g*) obtained using TSABC. Each of the 25 curves corresponds to a dataset simulated under Model C (left) and Model Gb (right) with no sequencing error (*ϵ* = 0), sample size *m* = 200 and sequence length **l** = 10^6^.

Table 3 shows that TSABC performs similarly to the results reported by [18] despite a 4 000-fold reduction in data used for inference, and despite the challenges we imposed on TSABC: gene conversion was incorporated in data simulation models but not the ABC inference simulations, and the latter also used a misspecified demographic model.

**Table 3.** Comparison of TSABC inference of *µ* (in units of 10^*−*8^) with results reported in [18]. TSABC results are obtained from 25 simulated datasets under each model.

|  | Model C |  | Model EA |  |  |  |
| --- | --- | --- | --- | --- | --- | --- |
| | $\hat{\mu}$ | SD | $\hat{\mu}$ | SD | $m$ | $\ell$ |
| [18] | 1.30 | 0.020 | 1.34 | 0.007 | 4 000 | $10^8$ |
| TSABC | 1.30 | 0.017 | 1.28 | 0.007 | 10 | $10^7$ |

### 1000 Genomes data analysis

The global mean 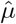 over the two chromosomes and eight populations is 1.27 *×* 10^*−*8^ (Table 4), similar to previous estimates assuming *µ* to be constant over populations [26, 29, 30], and also those finding small between-family differences in *µ* [31, 32]. A two-way ANOVA revealed no significant difference between the two chromosomes, but highly significant differences across populations, which may be due to differences in heritable factors or environmental exposures.

**Table 4.** Estimates of the posterior mean and SE of the mutation rate per site per generation (in units of 10^*−*8^) on human chromosome 20 and 21 for populations MSL (Mende in Sierra Leone), LWK (Luhya in Webuye, Kenya), BEB (Bengali from Bangladesh), ITU (Indian Telugu from the UK), FIN (Finnish in Finland), GBR (British in England and Scotland), JPT (Japanese in Tokyo, Japan), and CHB (Han Chinese in Beijing, China). The TSABC analysis assumes the 1KGP demographic model in each population.

|  |  | MSL | LWK | BEB | ITU | FIN | GBR | JPT | CHB |
| --- | --- | --- | --- | --- | --- | --- | --- | --- | --- |
|  | Sample size: | 170 | 198 | 172 | 204 | 198 | 182 | 208 | 206 |
| Chr 20 | $\hat{\mu}$ | 1.27 | 1.24 | 1.23 | 1.22 | 1.32 | 1.36 | 1.32 | 1.20 |
|  | SE | 0.004 | 0.004 | 0.004 | 0.004 | 0.004 | 0.011 | 0.004 | 0.004 |
| Chr 21 | $\hat{\mu}$ | 1.25 | 1.26 | 1.21 | 1.29 | 1.29 | 1.35 | 1.33 | 1.24 |
|  | SE | 0.004 | 0.005 | 0.006 | 0.005 | 0.006 | 0.006 | 0.006 | 0.006 |
| Combined | $\hat{\mu}$ | 1.26 | 1.25 | 1.22 | 1.25 | 1.31 | 1.36 | 1.33 | 1.22 |
|  | SE | 0.003 | 0.003 | 0.003 | 0.004 | 0.004 | 0.006 | 0.004 | 0.004 |

Figure 8 shows positive growth in the past 1 000 generations for all eight populations. CHB and BEB (both in Asia) have the highest *N* (0) while MSL and LWK (both in Africa) have the lowest *N* (0) despite having the highest values of *N* (1000). These findings are consistent with the results of [26] for 400 *≤ g ≤* 1 000, and the recent growth estimates obtained using Relate [11, Figure 3].

**Fig 8.**
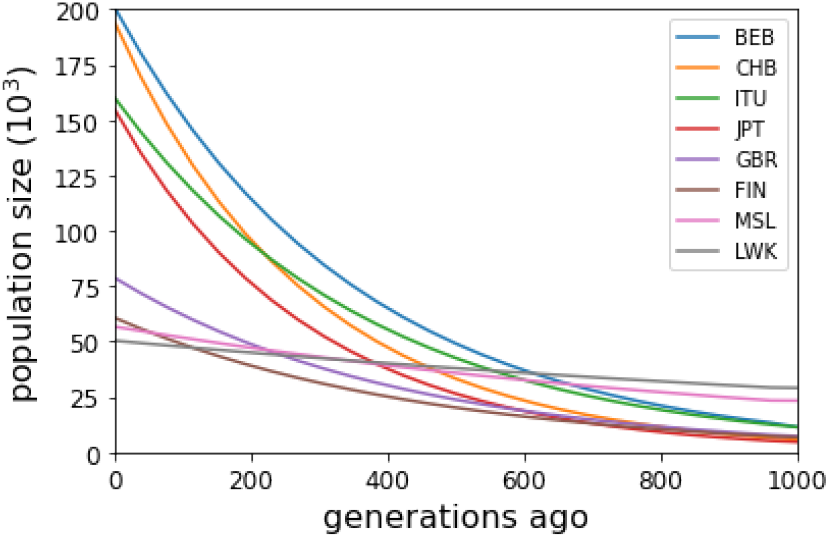
Estimates of recent population sizes for eight populations sampled in the 1000 Genomes Project (curves are shown in order of decreasing *N* (0)). See Table 4 caption for explanation of the population labels.

## Discussion

We have shown that ARG-derived IBD combined with ABC can deliver big advantages over previous IBD-based methods for inferring evolutionary and demographic parameters from a sample of genome-wide sequences, including nonparametric estimation of past population sizes. Despite verifying that IBD extracted from an inferred TS is often inaccurate, we have shown that it provides powerful inferences for the mutation rate and historic population sizes. For example, we obtained similar estimation results to a previous study that used 4 000 times more data for inference. These advantages arise because we can define IBD in terms of a common MRCA, avoiding both the problem of detecting recombinations and the need for a minimum IBD length. Further, we require only IBDs from *m* sequence pairs, rather than all *m*(*m−* 1)*/*2 pairs, which reduces computational effort with little loss of statistical efficiency.

We illustrated our TSABC approach in simple scenarios, finding that it suffers only modest loss of efficiency relative to using true IBD. Importantly, removing IBDs with length below even a low threshold reduces the precision of inferences despite the poor quality of ARG-based IBD inferences.

TSABC can be computationally demanding for complex demographic models, and the results presented here are limited to inferring the mutation rate and two parameters of a demographic model. However, we were able to incorporate unknown nuisance parameters such as the sequence error rate and misspecification of the demographic model to challenge TSABC inference without substantial detriment to inference quality.

Our results open the way for more powerful demographic and evolutionary inferences from samples of genome sequences than have previously been available.

### Data availability

Data and code used here are available at:

github.com/ZhendongHuang/Estimating_evolutionary_and_demographic_parameters_Huang

## Acknowledgment

ZH is funded by Australian Research Council grant DP210102168 awarded to YBC, DJB and JK. JK acknowledges support from the Robertson Foundation, US National Institutes of Health (grants HG011395 and HG012473) and UK Engineering and Physical Sciences Research Council (grant EP/X024881/1).

## Supporting Information

### S1 Appendix: The efficient IBD subsets and TSABC algorithms

Algorithm 1 searches the IBDs in the order of their children. For instance, the algorithm first starts to search all IBDs corresponding to the sequence pair (1, 2) along the whole sequence, from left to right. Then it searches the IBDs for (2, 3) and so on until the final sequence pair (*m*, 1).

Algorithm 2 adopts a standard ABC approach, the innovation here is in the choice of summary statistics which are derived from an inferred TS.

### S2 Appendix: Derivation of Estimators when TS is known

#### S2.1 Mutation and sequencing error rates

The right endpoint *r*_*i*_ of an IBD segment is usually the site of an effective recombination event for (*c*_1_, *c*_2_), where “effective” means that *c*_1_ and *c*_2_ have a different MRCA on either side. The exceptions are IBDs terminating at the sequence end site **l**, which are excluded from the derivations below but, as they are rare, there is little impact if they are included in practice. For the derivation of our estimators, we also assume that at most one mutation occurs at each site since the MRCA of the sequences at that site. The effect of this assumption is minor when the mutation rates is low, which is the case for humans and many other organisms. We further assume that sequencing errors occur independently at rate *ϵ* at each site, and do not occur at sites with mutations.

##### Algorithm 1 The efficient IBD subsets algorithm

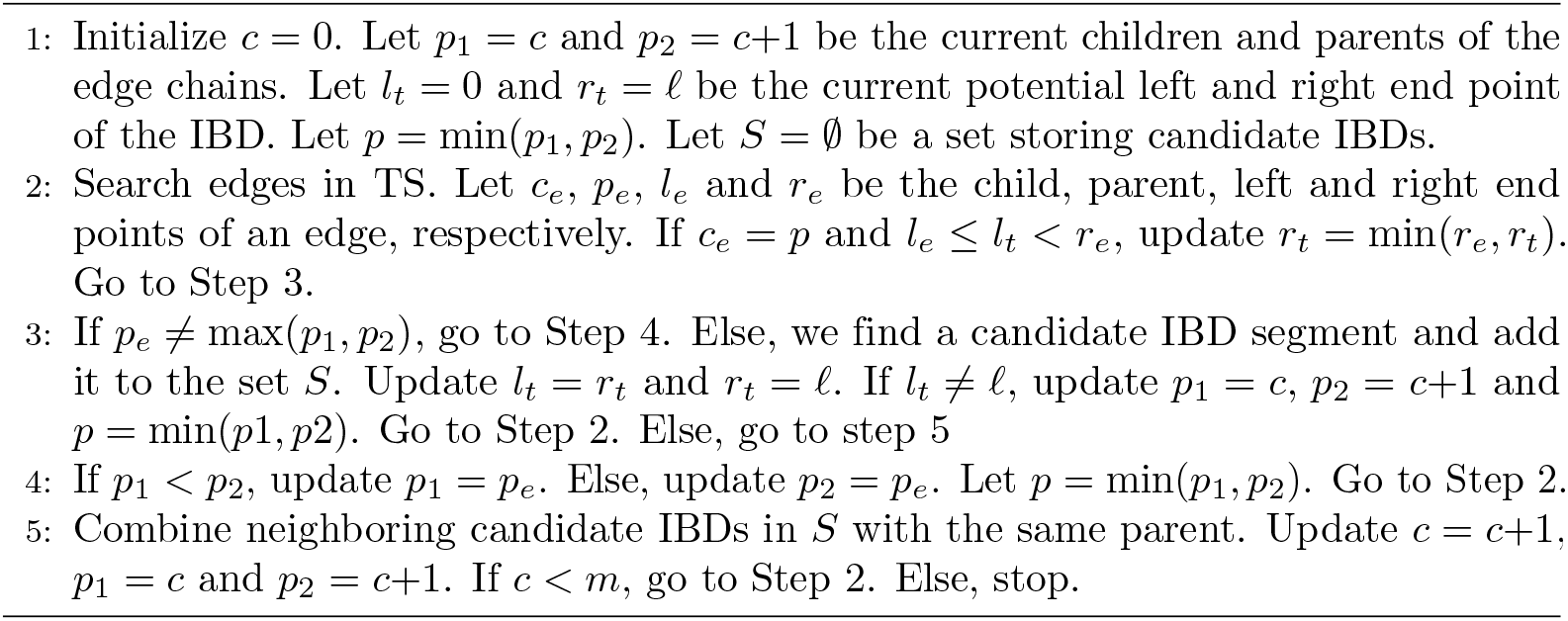

##### Algorithm 2 TSABC algorithm for parameter *θ*

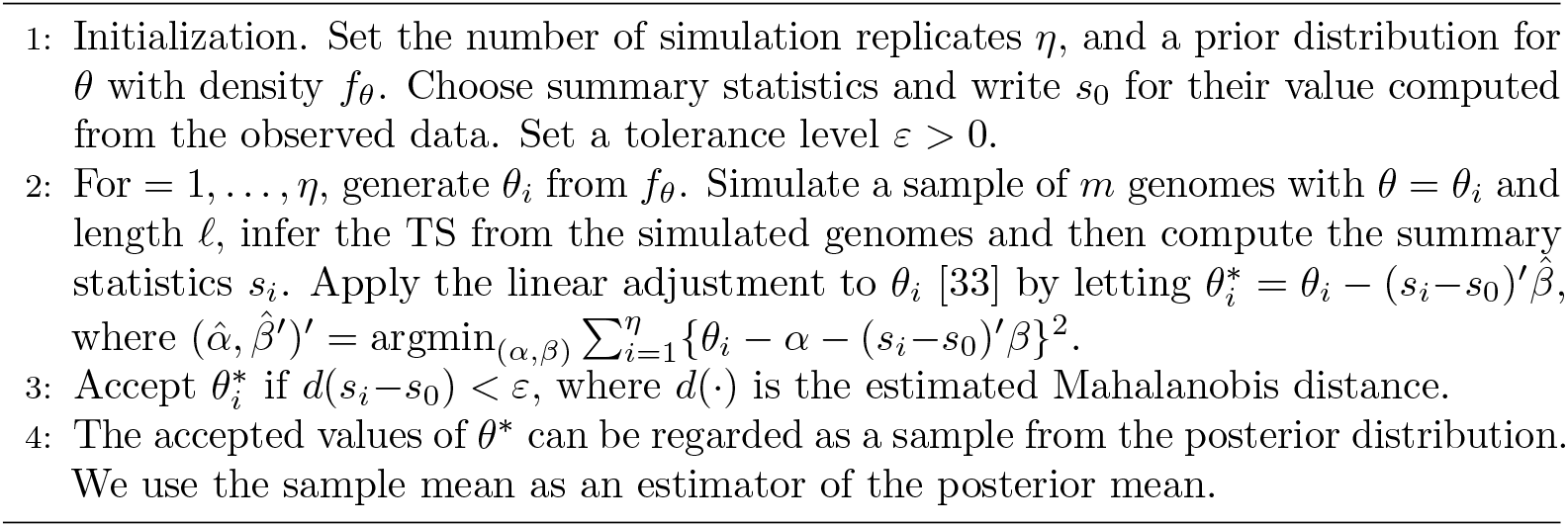

Suppose that a recombination occurs at site *s* of sequence *c*_1_ creating two subsequences going backward in time, in the intervals [1, *s*] and [*s*+1, **l**]. This recombination is effective for (*c*_1_, *c*_2_) if and only if one of these subsequences coalesces (reaches a common ancestor) with *c*_2_ before coalescing with the other subsequence. By symmetry, all three possible coalescence events are equally likely, and so the recombination has probability 2*/*3 of being effective. Mutation and effective recombinations occur independently at each site of *c*_1_ and *c*_2_. Given that one of these events occurs at a site before the two sequences reach their MRCA, with probability (2*r/*3)*/*(*µ*+2*r/*3) it is an effective recombination. Thus the expected number of mutations that occur in the segment before it is terminated by an effective recombination is one less than the mean of a geometric distribution with parameter (2*r/*3)*/*(*µ*+2*r/*3), which is 3*µ/*2*r*. Site differences in IBDs can also arise from sequencing errors, which occur with rate 2*ϵ* per site.

Let 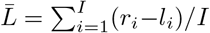 and 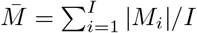denote the averages of the IBD segment length and the number of sites that differ in an IBD segment, respectively. Our first estimating equation is

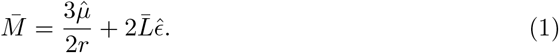

which can be read intuitively as site differences = mutations + sequencing errors. Our second estimating equation has a similar interpretation, but is based on site differences on each sequence relative to both its neighbours in the efficient subset, rather than between pairs of sequences:

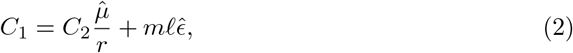

where we define

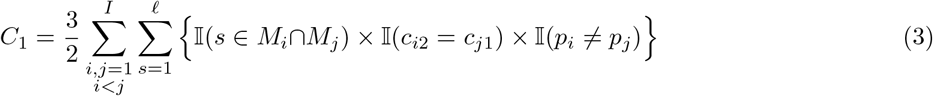

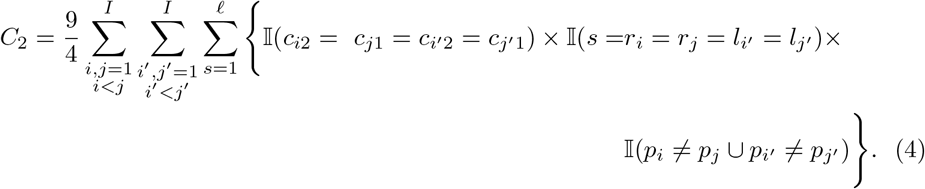

These quantities estimate, respectively, the total number of sequencing errors and mutations (*C*_1_) and the number of recombinations (*C*_2_) on the branch immediately above sequence *c*, before the first coalescence between any of *{c−* 1, *c, c*+1*}*. See below for further explanation. Among all of the target recombinations, only 4*/*9 of them can be unambiguously determined from the data, so we scale this count by 9*/*4 in (4). The factor 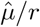 in (2) converts the estimated number of recombinations to an estimated number of mutations. The final term in (2) is the expected total number of sequencing errors among the *m* sequences, each of length **l**.

Combining (1) and (2), we obtain

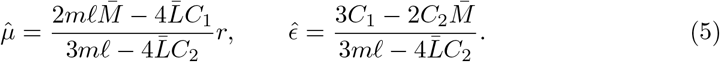

#### S2.2 Time since MRCA of IBDs

Given *g*_*i*_, the coalescence time of *c*_*i*1_ and *c*_*i*2_, the probability of a recombination event being effective (and thus being the right end point of the IBD) is no longer 2/3, but a function of *g*_*i*_. For example, if a recombination event occurs more recently than the coalescence, when *g*_*i*_ is small there is little opportunity for the two subsequences created by the recombination to find a common ancestor before *g*_*i*_, which implies a high probability for this recombination event to be effective. For this reason, it is difficult to estimate *g*_*i*_ based on the distribution of the recombination events or IBD lengths.

Note that |*M*_*i*_| follows a Poisson distribution with parameter 2(*µg*_*i*_ + *ϵ*)(*r*_*i*_ *− l*_*i*_).

Given 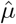,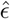, we find the first moment estimator of *g*_*i*_ by solving, for *p ∈ P*,

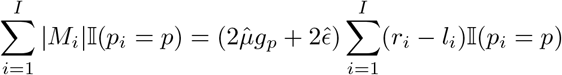

to obtain

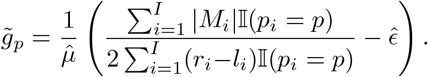

Let *g*_0_ = 0, and noting that *g*_*p−*1_ *< g*_*p*_ for *p* = *m*+2,…, *n*, we estimate *g*_*p*_ by the following quadratic optimization with linear constraints,

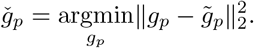

subject to *g*_*p*_′ − *g*_*p*_ ≥ 0, for *p* ′ > *p* and *p* ′, *p ∈ P*.

By the nature of constrained optimization, many parent nodes will be inferred to share the same age, which is unrealistic. Numerical studies show that it will be helpful to smooth these estimates when they are used later in estimating the population size. For this reason, the final estimate ĝ_*p*_ of *g*_*p*_ is acquired by further smoothing 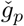 by a Savitzky-Golay smoothing filter [34].

#### S2.3 Present and past population sizes

We estimate population sizes by first estimating the density 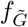 of 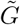, the TMRCA at a specific site *s*. We estimate 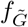 by relating it to the density *f*_*G*_ of *G*, the TMRCA of an IBD segment. We first estimate *f*_*G*_ empirically from the estimated TMRCAs of each IBD segment, as calculated in Section S2.2. Then the density of 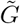 is derived by conditioning on *L*, the length of the IBD segment. Intuitively, the larger *L* is, the more sites are covered by the IBD. Hence

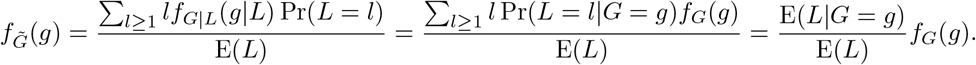

The mean IBD length E(*L*) can be estimated as 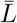, and the conditional mean E(*L*|*G*) can be found by a local linear kernel regression estimator given each pair 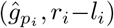 of IBD_*i*_, where 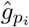 is the estimated TMRCA in Section S2.2. Thus, the estimate 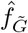 of 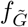 can be found by substituting the corresponding estimates of *f*_*G*_, E(*L*| *G*) and E(*L*). We then smooth the estimate 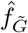 by a Savitzky–Golay filter.

To estimate population sizes, note that the distribution of 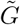 is solely determined by the coalescent rate 1*/N* (*g*), i.e.,

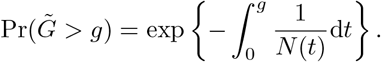

Taking the log-derivative with respect to *g* on both sides, we have

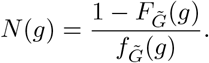

We thus calculate the estimate

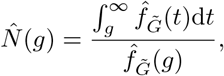

followed by another Savitzky–Golay smoothing filter.

#### S2.4 Interpretation of *C*_1_ and *C*_2_

The quantity *C*_1_ estimates the total number of sequencing errors and mutations on the branch immediately above (i.e. backwards in time from) a sequence *c*, before the first coalescence between any of *c, c −* 1 and *c*+1.

In the efficient IBD subset, for each sequence *c* we record the IBDs of the pairs (*c−* 1, *c*) and (*c, c*+1). If IBDs in (*c−* 1, *c*) and (*c, c*+1) covering a site *s* have different parent nodes, then *c* must coalesce with either *c−* 1 or *c*+1 at site *s* more recently than the coalescence of *c−* 1 with *c*+1. In this case, any site differences contained in both IBDs can be attributed to an event unambiguously located on the branch immediately above sequence *c* before the first coalescence.

We also wish to include site changes above *c* in the case where *c −* 1 coalesces with *c*+1 first, in which site differences cannot be located in this way. By symmetry, this case occurs 1*/*3 of the time, and so we scale the previous count by a factor of 3*/*2. See Figure S1 (top left) for illustration. The quantity *C*_1_ in (3) thus sums the total number of sequencing errors and mutations on the branch immediately above each sequence *c*, before the first coalescence.

Likewise, the quantity *C*_2_ estimates the total number of recombinations on the branch immediately above each sequence *c*, before the first coalescence between any of *c, c−*1 and *c*+1. Recall that *p*_*i*_, *l*_*i*_ and *r*_*i*_ are the MRCA and left and right endpoints of IBD_*i*_. Similarly to *C*_1_, we only count recombinations that produce four adjacent 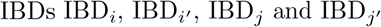and 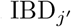, with the first two corresponding to sequence pair (*c−*1, *c*) and the latter two corresponding to (*c, c*+1), such that 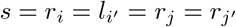 (i.e., the breakpoint between *i* and *i*^*′*^ is the same as the breakpoint between *j* and *j*^*′*^), and either *p*_*i*_≠*p*_*j*_ or 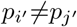, as shown in Figure S1 (top right). We then scale to account for the remaining cases.

If we have two IBD breakpoints at *s*, we must have a recombination on the *c* lineage since we assume that only one recombination can occur at *s*. If the MRCAs of these IBDs also satisfy *p*_*i*_≠*p*_*j*_ or 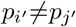, the recombination must occur before any coalescence, since:

- if *{c−*1, *c*+1*}* coalesce before the recombination, we must observe *p*_*i*_ = *p*_*j*_ and 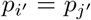;
- if *{c−*1, *c}* coalesce before the recombination, there would not be an IBD breakpoint at *s* for the (*c−*1, *c*) pair;
- likewise for when *{c, c*+1*}* coalesce before the recombination.

Thus we do indeed count a subset of the desired recombinations.

When there is a recombination in *c* before any coalescences, there are 4 lineages immediately after the recombination (backwards in time), as shown in Figure S1 (bottom). There are three cases:

- If lineages 1 and 2 coalesce first, the recombination is not effective and there are no IBD breakpoints at *s* (probability 1*/*6).
- If lineages 3 and 4 coalesce first, we will have 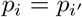 and 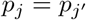 and so not count the recombination (probability 1*/*6).
- Otherwise, we may count the recombination (probability 2*/*3).

As shown in Figure S1 (bottom), suppose (without loss of generality) that for the third case, lineages 1 and 3 coalesce first. If the coalesced lineage then coalesces with lineage 2, then there will not be an IBD breakpoint at *s* for (*c, c*+1); otherwise the required pattern will be produced. Thus we only count 2*/*3*×* 2*/*3 = 4*/*9 of the cases, and so scale by a factor of 9/4 to estimate the desired number of recombinations.

### S3 Appendix: Further details for 1KGP data analysis

The chromosome lengths are **l**_20_ = 63 025 522 and **l**_21_ = 48 129 897 sites, of which 1 552 394 and 927 753 sites are polymorphic in the full dataset. The sequence data were downloaded as .vcf files from ftp.1000genomes.ebi.ac.uk. Then, we converted them to the .samples format required for input to tsinfer and adopted human reference assembly GRCh37 recombination map following the data pre-processing steps in [35]. Specifically, we first cloned the Github repository from github.com/awohns/unified genealogy paper and installed all of the necessary software, packages and modules listed in the “requirements.txt” file and the “tools” sub-folder. Then we redirected to the “all-data” sub-folder and conducted the “Makefile” document to build the tree sequence for 1000 Genomes chromosome 20, during which the program downloaded the chromosome 20 variant data and produced a .samples file (tsinfer input format) converted from a .vcf file. Then IBDs were extracted from the inferred TS, and TSABC was employed, as described in Section. A similar process is repeated for chromosome 21.

**Fig S1.**
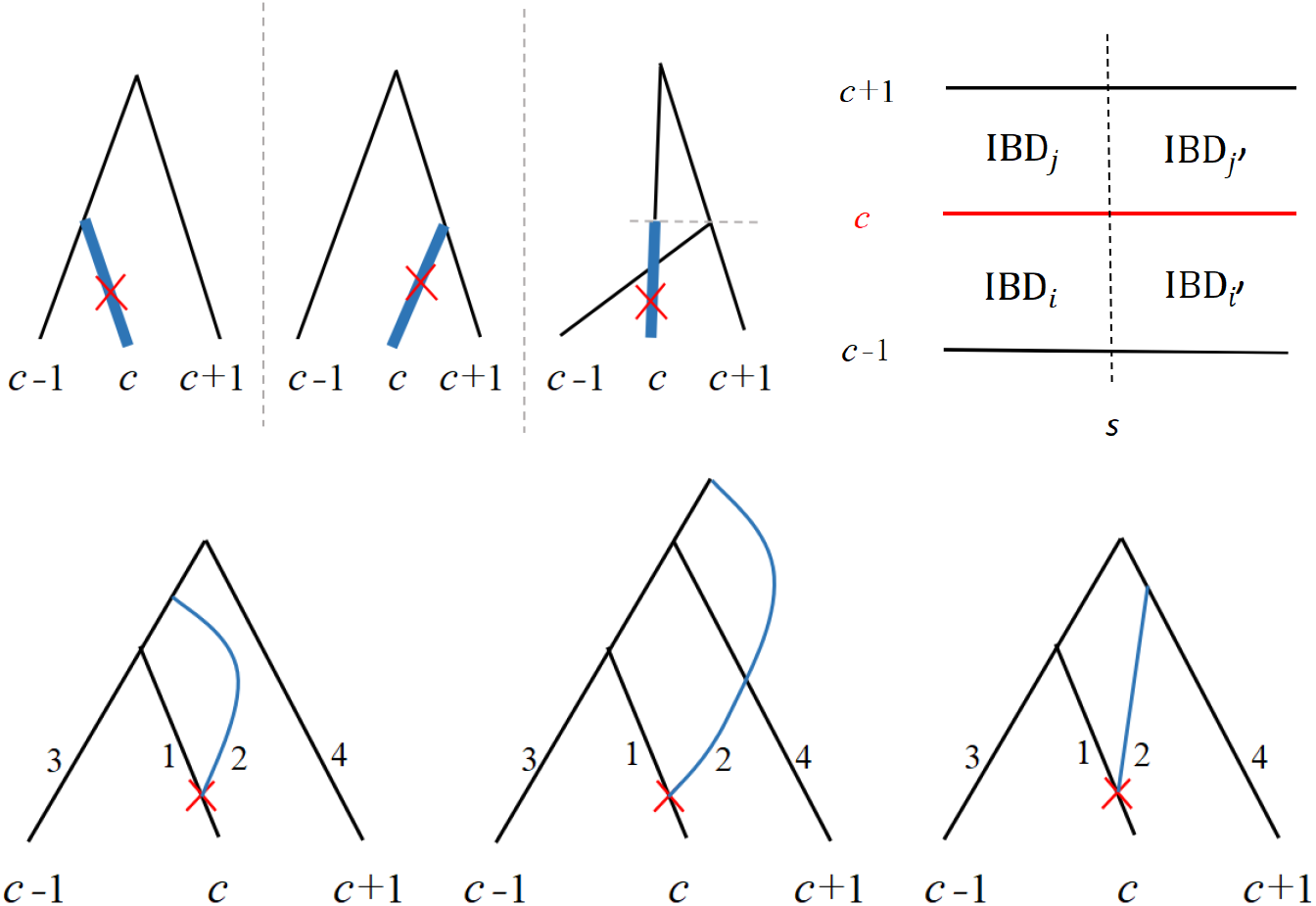
Top left: the three possible coalescent patterns of sequences *c, c−*1 and *c*+1 at a site *s*. While mutation events on the thicker edges should be included in the quantity *C*_1_, only those in the first two patterns are counted. Top right: a sketch of four IBDs corresponding to sequence pairs (*c−* 1, *c*) and (*c, c*+1). Bottom: a recombination event occurred on sequence *c*, which breaks the sequence before any coalescence between *c−* 1, *c* and *c*+1, immediately resulting in a total of four segments (1, 2, 3, 4). The figure shows three of the possible coalescent patterns, corresponding to the cases where segments 1 and 3 coalesce first.

### S4 Supplementary Figures

**Fig S2.**
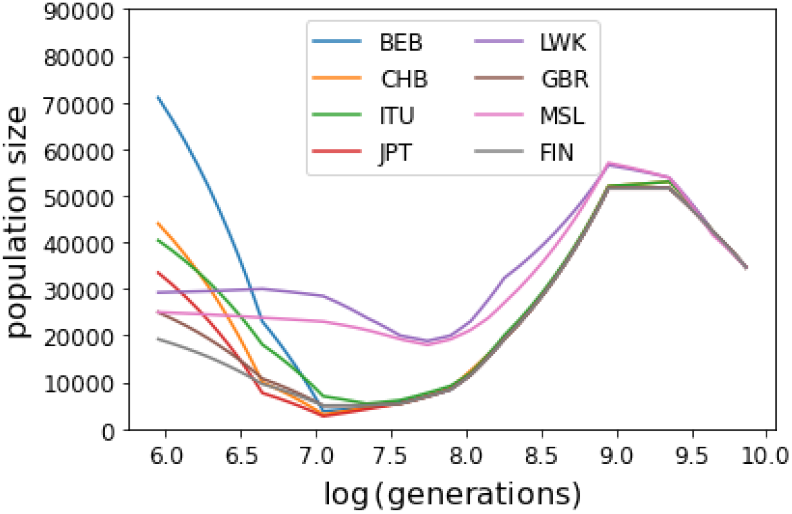
The 1KGP *N* (*g*) models [26]. Natural logarithms of *g* are shown on the *x*-axis, with the models starting at *g* = exp(6) *≈* 400 generations in the past. The values of *N* (1 000) which form the right endpoints of Figure 8 correspond to *x* = log(1 000) *≈* 6.9.

**Fig S3.**
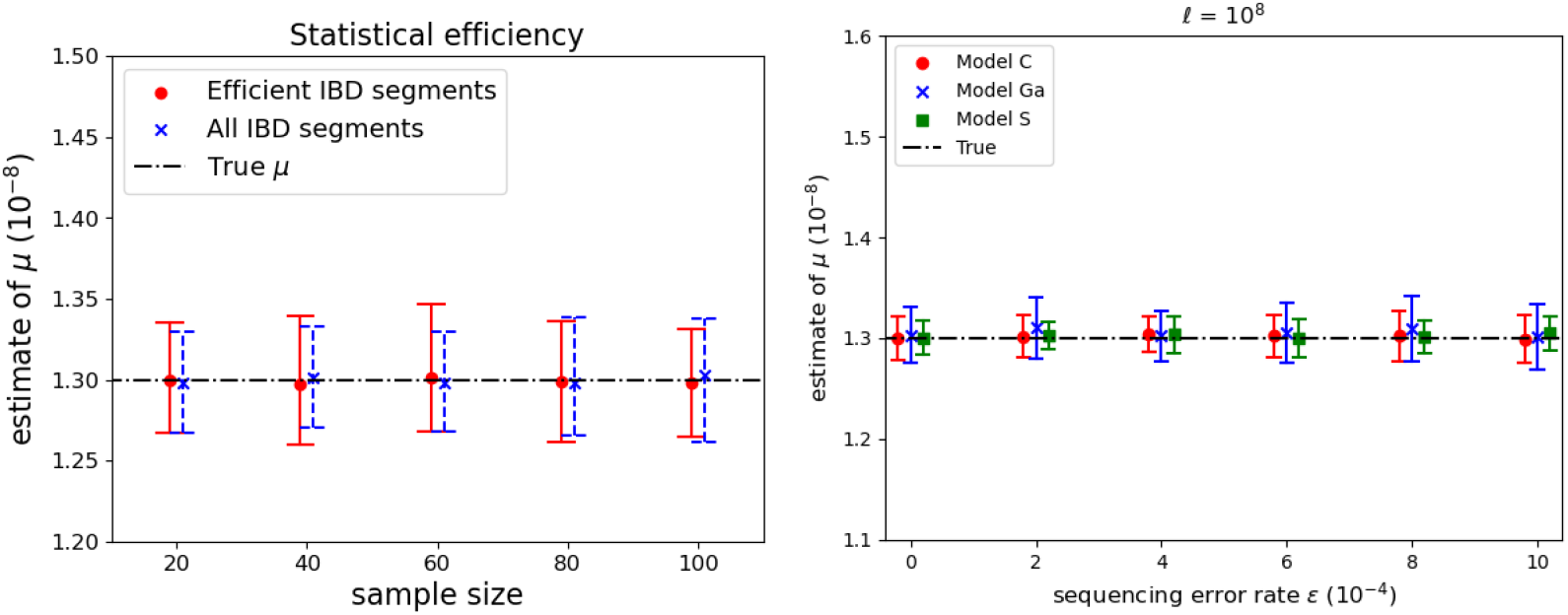
Left: Estimated 95% CIs for the estimation of *µ* when an efficient subset of IBD segments were extracted from the TS and when all IBDs were used. At each sample size, 25 replicate datasets were simulated under Model C, with sequence length **l** = 10^7^. Right: Impact of sequencing error rate *ϵ* on 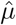 under Model C, Model Ga and Model S, from 25 replicates with sample size *m* = 10 and sequence length **l** = 10^8^.

**Fig S4.**
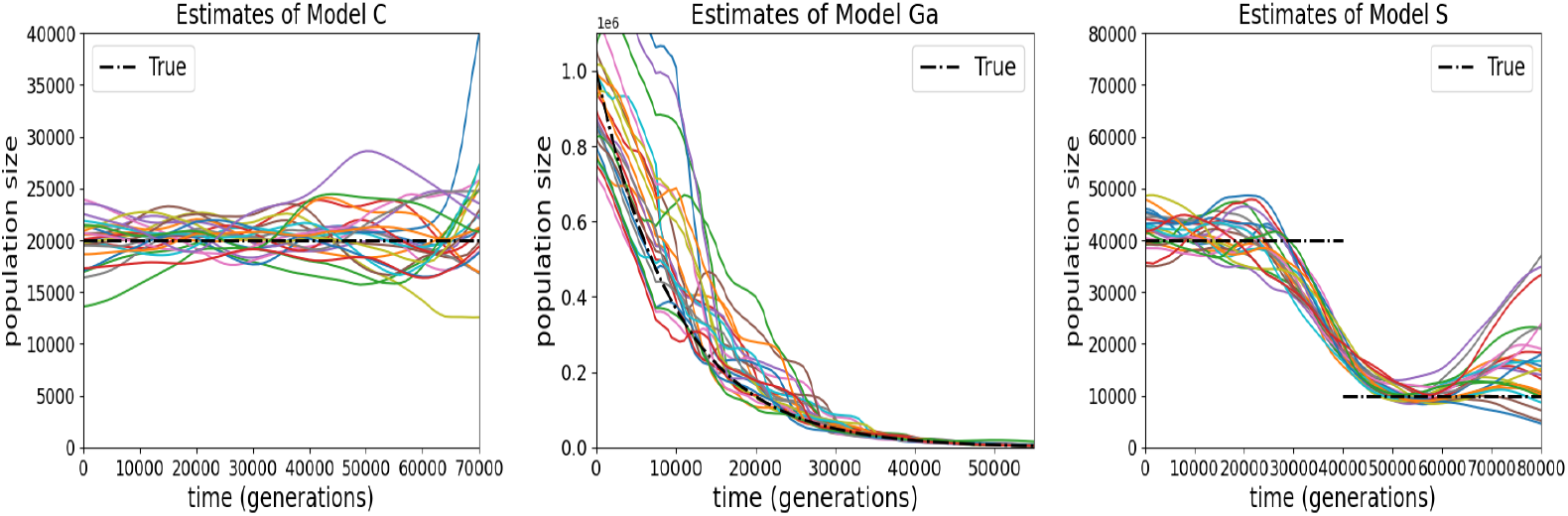
Estimates of the population size *N* (*g*) under Model C, Model Ga and Model S, from 25 simulations at each setting. Sequence length **l** = 10^7^, sequencing error rate *ϵ* = 10^*−*3^, sample size *m* = 80.

